# Cover crops improve early season natural enemy recruitment and pest management in cotton production

**DOI:** 10.1101/786509

**Authors:** Carson Bowers, Michael Toews, Yangxuan Liu, Jason M. Schmidt

## Abstract

A shift to more ecologically based farming practices would improve the sustainability and economic stability of agricultural systems. Habitat management in and around agricultural fields can provide stable environments that aid in the proliferation of natural enemy communities that moderate pest populations and injury. Winter cover crops offer a potentially cost-effective approach to improving habitat that supports natural enemy communities early in the growing season. We investigated the effects of winter cover crops including cereal rye (*Secale cereal* L.) and crimson clover (*Trifolium incarnatum* L.) on the abundance and diversity of natural enemies, key pest populations, biological control services, and cotton yield. Winter cover crops were established on 0.4 ha replicated field plots in the fall of 2017 and 2018. Suction sampling during each cotton development stage demonstrated that a rye cover crop promoted greater abundance and diversity of natural enemy communities in early cotton stages. Extensive leaf sampling of seedling cotton showed that cover crops significantly reduced thrips infestations. Furthermore, stink bug boll injury decreased on plots prepared with a rye cover compared to cotton lacking this additional habitat. Combining end of season yield results and management practices with an economic analysis of the costs of production, the value of cotton grown into a cover crop was cost competitive with conventional (no cover) cotton production. These results suggest that conventional growers utilizing cover crops could reduce insecticide inputs through natural reductions in pest pressure, and overall do not incur additional production costs.

## 1. Introduction

Meeting food and fiber demands of a growing population is often focused on increasingly intensive agricultural practices including synthetic fertilizers and pesticide applications (Matson et al., 1997; Fedoroff et al., 2010; Tilman et al., 2011). Although these intensive practices can support high levels of production, there are recognized negative environmental trade-offs to biodiversity (Tscharntke et al., 2005; Landis, 2017; Wittwer et al., 2017), water quality (Matson et al., 1997), and soil health (Stavi & Lal, 2015). Balancing reliance on synthetic inputs with biologically based, ecosystem services (e.g. biological control, pollination, nutrient cycling), is advocated to promote environmentally friendly solutions for the challenge of feeding a growing population (Rusch et al., 2017; Wittwer et al., 2017; Murrell, 2017; Kaye & Quemada, 2017; Tilman et al., 2011; Landis, 2017). This shift, known as ecological intensification, can improve the long-term sustainability and ecological stability of agricultural ecosystems.

Promoting ecosystem services can be achieved through improving conservation practices that favor local biodiversity combined with reduction in broad spectrum insecticides (Shields et al., 2018; Snyder, 2019, Gurr et al., 2017; Begg et al., 2017; Gurr et al., 2000, Landis et al., 2000). For example, managing the habitat in and around agricultural fields can mitigate some effects of intensive management to increase the effectiveness of natural enemies by promoting more abundant and diverse predator communities (Gurr et al., 2017). A variety of mechanisms are proposed for the effects of habitat management on predator communities, including: provisioning of alternative prey, and minimizing intraguild predation via additional microhabitat availability (Finke & Denno, 2002; Janssen et al., 2007). Habitat fueling alternative prey availability helps sustain natural enemy populations during periods when there is little to no crop habitat (Staudacher et al., 2018; Gardarin et al., 2018; Roubinet et al., 2017). Thus, efforts to promote habitat diversity may create a “resource bridge” to build diverse natural enemy communities ready for invading pests.

Winter cover crops are a form of habitat management that adds habitat complexity into cropping systems, and enhances multiple ecosystem services critical to sustainable crop production (Daryanto et al., 2018; Duzy & Kornecki, 2017; Adhikari et al., 2017). Cover crops are often planted to suppress weeds and reduce erosion, and provide several additional benefits such as improving soil health by fixing nitrogen, sequestering excess soil nutrients (Hartwig & Ammon, 2002), or sequestering atmospheric carbon and building soil organic matter (Kaye & Quemada, 2017). Additionally, cover crops and in field cover crop residue provide habitat diversity, which may improve natural pest control services by increasing the abundance and effectiveness of natural enemies in field.

Winter cover crops often benefit natural enemy communities and pest control in perennial systems (Gomez et al., 2018; Dong et al., 2018; Vogelweith & Thiery, 2017; Burgio et al., 2016). Yet, the response of natural enemies to cover crops and other habitat management strategies in row crop systems is variable (Daryanto et al., 2018; Begg et al., 2017; Tscharntke et al., 2016). In annual row crop systems, positive effects of cover crops on natural pest management appear driven by a combination of cropping system, geographical region, and cover crop type (Hooks et al., 2013; Mollot et al., 2012; Koch et al., 2015; Manandhar et al., 2017). Hence, there is a need for system and region specific studies examining the impact of cover crops on natural enemy communities and pest complexes. Additionally, few studies link changes in predator abundance and diversity to increased biological control services (Furlong & Zalucki, 2010), and for cover cropping to be fully integrated into the pest control tool box, any benefits must be linked to production value (Gurr et al., 2017).

Current insect pest management programs and pesticide usage for cotton in the southeastern United States are focused on early season thrips and a complex of late season stink bugs (Tillman, 2012; Lahiri et al. 2018). Thrips in the genus *Frankliniella* are a serious pest of seedling cotton as well as several other crops worldwide (Toews et al., 2010; Greenberg et al. 2009; Mouden et al. 2017). In previous studies, strip tillage and cover crops helped suppress early season thrips populations (Toews et al., 2010; Knight et al. 2017; Manandhar et al., 2017). Conversely, stink bugs, a mid-to-late season cotton pest, are challenging to control and force producers to rely on broad spectrum insecticides such as pyrethroids and organophosphates (Roberts & Toews, 2015), which are harmful to natural enemy communities (Gurr et al., 2017; Isaacs et al., 2009). There is building interest in management practices that reduce chemical inputs in favor of more environmentally friendly and sustainable options.

In this study, we investigated the effects of winter cover crops including cereal rye (*Secale cereal* L.) and crimson clover (*Trifolium incarnatum* L.) on the abundance and diversity of natural enemies, key pest populations, biological control services, and cotton productivity. We hypothesized that (1) the presence of cover crop residue increases the abundance and diversity of natural enemies, (2) the presence of a cover crop reduces numbers of early season thrips and late season stink bug pests, (3) cover crops improve biological service delivery on sting bugs by indirectly increasing predation on stink bug eggs, and (4) cover crops have positive effects on end of season production.

## 2. Materials and Methods

### 2.1 Study Site and Experimental Design

We investigated the effects of cover crops in a Georgia cotton production system by establishing plots in the fall of 2016 and 2017 at the UGA Southeast Georgia Research and Education Center at Midville, GA (Burke County, 32°52’15.6“N 82°13’12.0”W). The experimental design consisted of 0.4 ha plots (roughly square) organized in a completely randomized block design (n=4/treatment) for a total of 12 plots each sample year. All plots were separated by 3.6 m rolled rye alleyways. A control (no cover) was maintained throughout the off-season and managed following conventional tillage and winter herbicide applications common to southeast cotton production (Supplementary Material, Table A.1). Crimson clover (27 kg/ha) and rye (67 kg/ha) cover crops were planted early November using a cultipacker or grain drill and chemically terminated and rolled using a straight bar roller crimper 14 days prior to cotton planting. Cotton was planted into cover treatments May 5, 2017 (PHY 490 W3FE) and April 28, 2018 (PHY 440 W3FE) using a Unverferth strip till rig leaving an 8-inch tilled strip to serve as a seed bed, while conventionally tilled plots were disked followed by a rip and bed pass. All fields were irrigated during cover crop growth and the cotton growing season, and received no insecticides throughout the study (for full plot management details see Supplementary Material, Table A.1).

### 2.2 Arthropod Sampling

Canopy and ground dwelling arthropods were sampled using a 27.2cc modified reverse-flow leaf blower (SH 86 C-E; Stihl, Waiblingen, Germany) containing an average air velocity of 63 m/s with a mesh bag over the intake to collect natural enemies within a 1 m^2^ area quadrat. These areas within each plot were delineated by placing a 1 m^2^ quadrat, custom fabricated from 0.48 cm thick clear acrylic sheet with 0.3 m tall walls and metal bottoms, on the soil surface to prevent escape by ground dwelling or low-flying arthropods. Actual sampling locations were randomly selected on each sample date and all samples were at minimum 10 m from the plot edge (3 samples per plot). All cotton plants and cover crop residue within the 1 m^2^ area was suctioned (∼1 min/sample) until there was no visual arthropod activity on the ground or in the canopy after visual inspection. Suction sampling occurred during each of the primary cotton development stages (pre-emergence, seedling, vegetative growth, squaring, flowering, boll development), for a total of 6 sampling dates per plot during the 2017 and 2018 cotton growing seasons (Supplementary Material, Table A.2). All samples were placed in plastic bags and immediately stored in on ice, and preserved at −20° C until identification. All arthropods were identified to the family level with adult natural enemies identified to genus and species, when possible.

Adult and nymph thrips (Thysanoptera: Thripidae) populations were assessed in every plot during seedling growth stages at 14 and 21 days after cotton planting following standard protocols (Toews et al., 2010). On each date, two samples of 5 plants were randomly collected in a diagonal transect (across rows) starting at least 10m from field margins. Briefly, whole cotton seedlings were removed from the soil and immediately inverted in 0.47 L glass jars (5 plants/jar) partially filled with 70% ethanol where the plants were vigorously agitated to dislodge thrips. In the laboratory, the alcohol was passed through a 125 μm sieve and the thrips were retained on the sieve and then gently washed onto gridded filter paper, identified and enumerated under a dissecting microscope.

Stink bug pressure was evaluated during cotton anthesis using sweep nets and boll injury assessments (following protocols by Toews et al., 2008). The primary stink bug complex in the southeast cotton cropping region is made up of the southern green stink bug, *Nezara viridula* L., the green stink bug, *Acrosternum hilare* Say, and the brown stink bug, *Euschistus servus* Say. Two sweep net samples (20 sweeps/transect) were performed in each plot (one at 20 m from one edge of the plot and at a parallel location on the other side of the plot) and the number of stink bugs was recorded. Stink bug species and life stage were recorded and the combined count of all non-predatory stink bugs was used for analysis (Supplemental Material, Table B.1). For evaluation of feeding damage on cotton bolls, 2.3 to 2.7 cm diameter bolls (Willrich et al., 2004) were collected from the same two transects in each plot and checked for symptoms of stink bug feeding (20 bolls/plot for 2017, 40/ plot for 2018). Internal boll injury was defined by the presence of callus growths (warts) or stained lint (Greene et al., 1999, Bundy et al., 2000), with bolls classified as injured or uninjured. Stink bug sweeps and boll collection starting at the 2^nd^ week of bloom were performed for 4 weeks, once per week during the cotton anthesis.

### 2.3 Estimating Biocontrol Services

To evaluate biocontrol services provided by natural enemies, egg predation and parasitism were estimated from sentinel egg masses. At the beginning of 2017 and 2018 cotton seasons, *Nezara viridula* colonies were established from field collected adults, and maintained to produce stink bug egg masses. Egg masses (3-5 day old) were collected from the colony one day prior to field deployment and stored in a refrigerator at 4-8 ° C. Colony egg masses laid on paper towels were cut to remove excess paper material and stapled to a 3.0 × 3.5 cm index card. Egg masses were affixed to 4 cotton plants per plot during the cotton flowering period for a total of 3 dates in 2017 and 4 dates in 2018. Egg masses were attached to plants on the underside of a main stem leaf at the first node above white flower. Plants were selected in a square (5 × 5 m) around the center of the plot, and marked with colored tape for easy detection and retrieval. After a 48-hour period, egg masses were collected and transported to the lab where the number of eggs missing or damaged was used to assess mean predation rates (percent egg removal). All egg masses were photographed before they were placed in the field and after collection. Remaining eggs were incubated and monitored for emergence of parasitoids to estimate egg parasitism rates.

### 2.4 Cover Crop Biomass and End of Season Yield/Fiber Quality

During the 2018 season, live cover crop biomass was estimated prior to cotton planting. All above ground cover crop biomass within a 0.3 m^2^ area was clipped at the soil surface and collected (excluding conventionally managed plots), and three samples were collected from each plot. Collected samples were stored in paper bags and dried for 48 hours in a gravity convection oven at approximately 60 °C and dry weight recorded for each sample.

End of season cotton yield and fiber quality were estimated for both years by harvesting along two 24 m transects per plot using a mechanized two row spindle picker. Following harvest, seed cotton was weighed and then ginned on a per sample basis at the University of Georgia Cotton Micro Gin in Tifton, Ga to determine lint and seed fractions. Samples of lint were submitted to the Cotton Program Classing Office in Macon, GA to determine any differences in fiber quality.

To estimate the economic benefits of adopting cover crops in cotton production, the net return per acre was compared for each of the three treatments of cover crops (*i*) for 2017 and 2018 (*t*). The gross revenue was calculated using the lint and cottonseed yields and their historical prices each year.

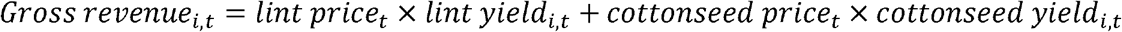

The prices for cotton lint include the cotton loan price and market price to compare the effect of prices on the profitability of different treatments. Cotton loan price is the minimum amount of money farmers would receive for their cotton with specific fiber quality. Market price for cotton lint were calculated from incorporating fiber quality by using the annual cotton price statistics from U.S. Department of Agriculture, Agricultural Marketing Service (USDA-NASS 2018). Costs were calculated based on the input uses and farming practices in planting cover crops, cultivating the land, planting cotton, herbicide applications, harvesting and ginning of cotton (for additional details of analysis see Supplementary Material, C.1). The cost was calculated using the following equation:

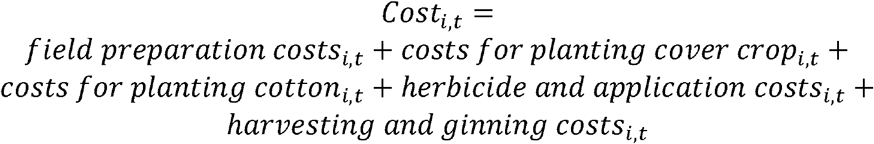

The net return per acre for cotton production was calculated for each of the treatment and each production season as follows:

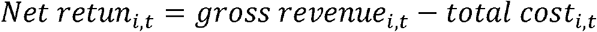

### 2.5 Data Analysis

Due to inter-annual variability, response variables were analyzed and displayed separately for 2017 and 2018. Predator density (number per m^2^) and Shannon diversity (H) were calculated for each sample throughout the growing season. Square root transformations for all count responses were required to normalize distributions and satisfy model adequacy. Predator density and diversity, stink bug and thrips abundance in relation to cover cropping treatments were analyzed using linear mixed effect models (LMM) (function = lmer, package = lme4; Bates et al., 2014) with date as a fixed effect, and plot as a random effect to account for repeated measures of plots over time. Production metrics were only estimated for the end of season, so a basic ANOVA model was used to compare production (i.e. yield, fiber quality, etc.; Table 1) between cover crop treatments. To test for significant effects of LMMs and ANOVAs, contrasts were evaluated at the 95% confidence interval with adjusted p-values for multiple comparisons using the “Tukey” method (function=lsmeans, contrast; package = lsmeans, Luke, 2017). Cumulative egg predation and proportion of injured bolls were evaluated in relation to cover crop treatments using generalized linear mixed-effect models (GLMM) for analysis of proportions (function = glmer, package = lme4) using plot and date as nested random effects (family=binomial). For significant effects of GLMMs, contrasts were evaluated at the 95% confidence interval with p-values adjusted using the “holm” method (function=glht, contrast; package = lsmeans, Luke, 2017). All statistical analyses were performed in R v 3.3.2 (R Core Team, 2018).

**Table 1.** Mean (±SE) end of season cotton production metrics for 2017 and 2018, by treatment. Results of ANOVA are shown. Different letters within rows indicate significant differences among means. Significant P values (α<0.05) are in bold. Fiber quality metrics used as response variables include lint yield (kg/ha), color grade, staple (length in 32nds of an inch), micronaire (mic), strength (grams/tex), reflectance (Rd), yellowness (+B), HVI length (inches), and uniformity. For full descriptions of fiber quality metrics see supplemental material (Table C.3).

| **2017** | **No cover** | **Crimson clover** | **Rye** | **F** | **df** | **P** |
| --- | --- | --- | --- | --- | --- | --- |
| *Lint Yield kg/ha* | 985.03(40.9) | 1005.78(70.87) | 1033.25(27.24) | 0.24 | 2,21 | 0.792 |
| *Color grade* | 29.75(1.25) | 26.00(1.89) | 27.25(1.83) | 1.29 | 2,21 | 0.296 |
| *Staple* | 35.75(.16) | 36.25(.25) | 36.38(.26) | 2.07 | 2,21 | 0.151 |
| *mic* | 4.15(.08) | 4.25(.05) | 4.35(.03) | 3.36 | 2,21 | 0.054 |
| *Strength* | 32.78(.36) | 33.05(.24) | 33.81 (.26) | 3.35 | 2,21 | 0.055 |
| *Rd* | 78.98(.27)<sup>a</sup> | 79.98(.26)<sup>b</sup> | 79.93(.33)<sup>b</sup> | 3.83 | 2,21 | **0.038** |
| *+B* | 8.03(.09) | 7.95(.08) | 7.78(.06) | 2.74 | 2,21 | 0.088 |
| *HVI Length* | 1.11(.00) | 1.13(.01) | 1.13(.01) | 2.30 | 2,21 | 0.125 |
| *Uniformity* | 82.88(.22)<sup>a</sup> | 83.23(.22)<sup>ab</sup> | 83.76 (.23)<sup>b</sup> | 4.01 | 2,21 | **0.033** |
| **2018** | **No cover** | **Crimson clover** | **Rye** | **F** | **df** | **P** |
| Live cover biomass(g) | NA | 2.50(0.69)<sup>a</sup> | 48.14(5.86)<sup>b</sup> | 59.83 | 1,22 | **0.001** |
| *Lint Yield kg/ha* | 929.49(48.07) | 1033.85(54.58) | 849.14(61.98) | 2.82 | 2,21 | 0.082 |
| *Color grade* | 36.38(2.03) | 34.88(1.89) | 34.75(1.83) | 0.22 | 2,21 | 0.803 |
| *Staple* | 37.63(.18)<sup>ab</sup> | 37.75(.25)<sup>a</sup> | 37.00(.19)<sup>b</sup> | 3.68 | 2,21 | **0.043** |
| *mic* | 3.71(.09) | 3.83(.09) | 4.00(.07) | 3.19 | 2,21 | 0.061 |
| *Strength* | 31.75(.28) | 31.99(.39) | 31.69(.39) | 0.19 | 2,21 | 0.825 |
| *Rd* | 74.39(.67) | 75.10(.92) | 76.36(.38) | 2.09 | 2,21 | 0.149 |
| *+B* | 8.78(.11)<sup>a</sup> | 8.63(.11)<sup>ab</sup> | 8.4(.07)<sup>b</sup> | 3.72 | 2,21 | **0.041** |
| *HVI Length* | 1.17(.01)<sup>ab</sup> | 1.19(.01)<sup>a</sup> | 1.16(.01)<sup>b</sup> | 3.53 | 2,21 | **0.048** |
| *Uniformity* | 81.24(.16) | 81.08(.29) | 81.86(.24) | 3.15 | 2,21 | 0.064 |

## 3. Results

### 3.1 Arthropod Sampling

#### 3.1.1 Natural enemies

A total of 2,675 predators were collected during the 2017 (1,123) and 2018 (1,552) sampling seasons (Supplementary Material, Table B.2). In 2017, predator density was significantly influenced by cover crop treatment (LMM: F_2,12_=3.90, *p*=0.049) and sampling date (LMM: F_5,216_=33.15, *p*<0.0001) with a significant interaction between cover crop treatment and date (LMM: F_10, 216_=8.41; *p*<0.0001; Fig. 1A). Likewise, in 2018 predator density was significantly influenced by treatment (LMM: F_2,217_=46.32, *p*<0.0001), date (LMM: F_5,217_=14.92, *p*<0.0001) and an interaction between treatment and sample date (LMM: F_10, 217_=10.30; *p*<0.0001; Fig. 1B). In the interaction between date and cover crop treatment indicates differences in predator density and diversity in the early growing season in both years in relation to cover crops (Fig. 1A and B; Supplemental Material, Table B.3). In 2017, a rye cover crop harbored significantly higher densities of natural enemies than no-cover conventional plots prior to seedling emergence (Fig. 1A). In 2018, a rye cover crop significantly improved predator density through seedling stage of cotton, while crimson clover showed no improvement over no-cover treatments (Fig. 1B; Supplemental Material, Table B.3).

**Fig. 1.**
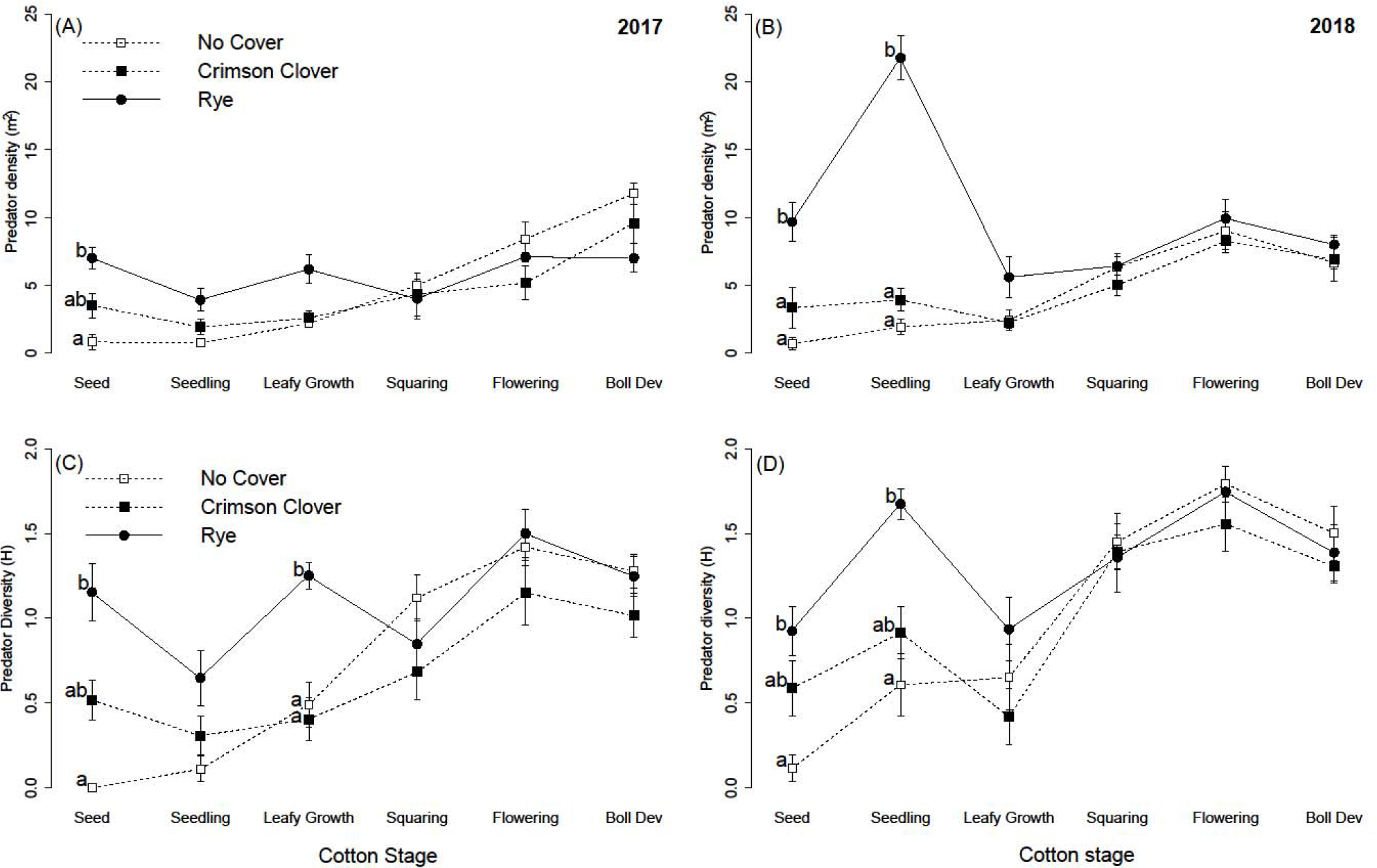
Predator density (no. predators per m^2^) (top panels A,B) and family level predator diversity (H) (bottom panels C,D) for the 2017 (left) and 2018 (right) sampling season across each major cotton development stage. Bars represent 1 SE of the mean for each treatment on each sample date. Letters indicate significant differences (α<0.05) among treatments on the same cotton development stage within years.

Similarly, we observed a significant effect of cover crop treatment (LMM: F_2,9_=8.58, *p*=0.0081), date (LMM: F_5,201_=26.32, *p*<0.0001) and a treatment by date interaction (LMM: F_10,201_=5.65; *p*<0.0001; Fig. 1C) in explaining diversity (H) of natural enemies in 2017. Similar effects of treatment (LMM: F_2,199_=8.69, *p*=0.0002), date (LMM: F_5,199_=26.22, *p*<0.0001) and treatment by date interaction (LMM: F_10,199_=3.32; *p*=0.0005; Fig. 1D) on natural enemy diversity were found in 2018. In 2017, the rye cover crop elevated predator diversity through the leafy growth stage of cotton compared to no-cover plots (Fig. 1C; Supplemental Material, Table B.3). In 2018, rye maintained significantly higher diversity of predators through the seedling stage compared to conventionally management plots (Supplemental Material, Table B.3).

#### 3.3.2 Thrips

Due to low thrips pressure, no assessment on thrips populations were feasible for the 2017 growing season (i.e. not detectible across all treatments). For 2018, the total number of thrips varied by sample date (LMM: F_1, 32_=122.7; *p*<0.0001), with a greater abundance of total thrips 21 days after planting (DAP) compared to 14 DAP (Fig. 2). The abundance of thrips adults was influenced by date (LMM: F_1,32_=120.33, *p*<0.0001), with a significant interaction between treatment and date (LMM: F_2, 32_=32.2; *p*=0.004). The interaction for adult thrips is explained by the fact that abundance was marginally affected by cover crop treatment 14 days after planting (Fig. 2A), though we found no difference in adult thrips abundance among treatments at 21 DAP. Nymphal thrips abundance was significantly influenced by treatment (LMM: F_1, 9_=5.54; *p*=0.027), date (LMM: F_1, 32_=21.05; *p*<0.0001) and an interaction between treatment and date (LMM: F_1, 32_=3.77; *p*=0.034). Rye and crimson clover cover treatments had significantly lower abundance of thrips nymphs on cotton seedlings compared to the no cover treatment 21 days after cotton planting (Fig. 2B), with no difference among treatments 14 days after planting.

**Fig. 2.**
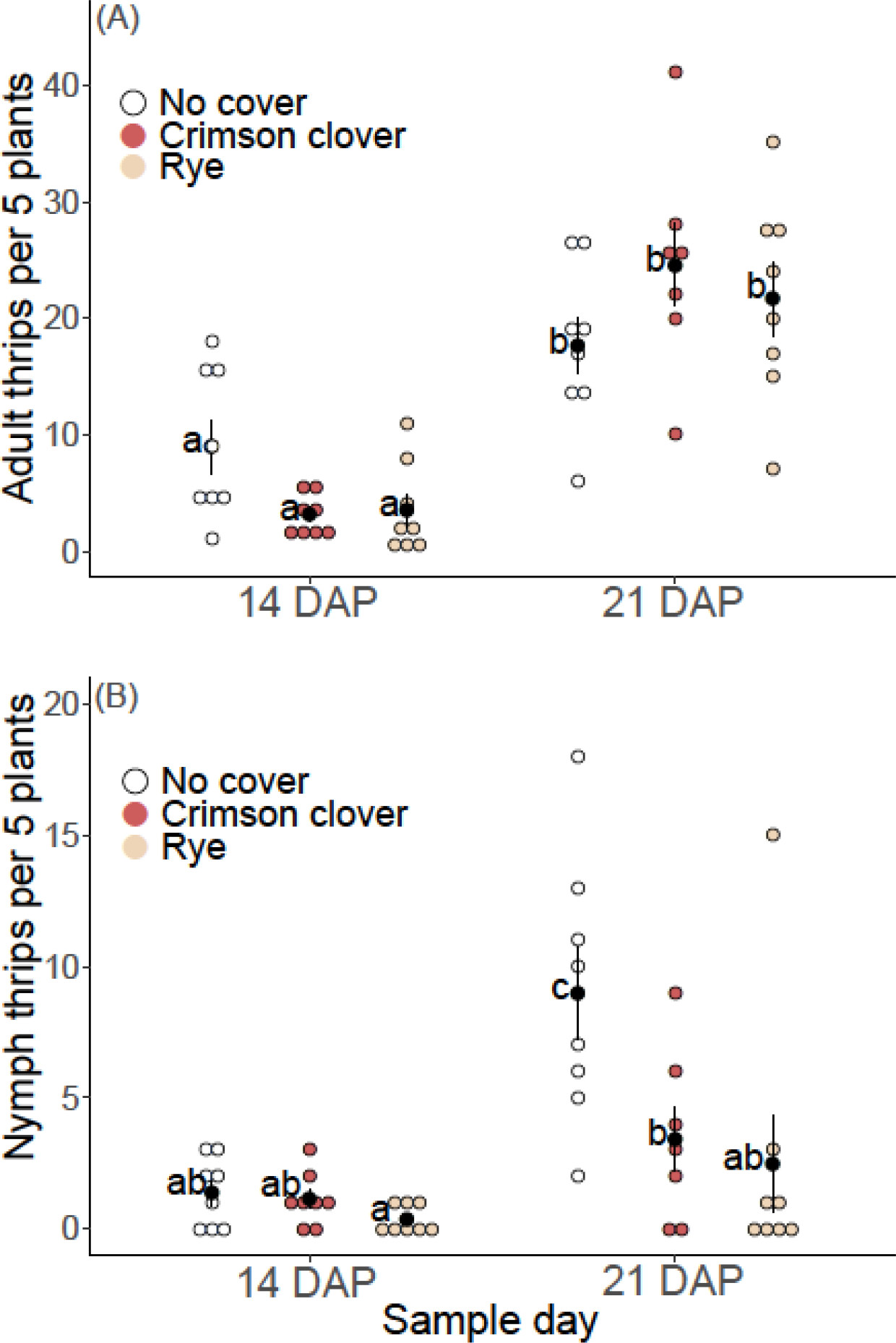
Number of adult (A) and nymph (B) thrips per 5 plants 14 and 21 days after cotton planting (DAP) in Rye, Crimson Clover, and No-cover treatments. Black dots and error bars represent mean and SE. Letters next to dots indicate significant differences (α<0.05) in thrips of the same life stage among treatments.

#### 3.1.3 Stink Bugs

The effect of cover crop treatments on stink bug abundance and boll injury was assessed during both sample years. The most common stink bug species collected in sweep samples was *N. viridula*, making up 84% of all stink bugs sampled in 2017, and 38% in 2018 (Supplementary Material, Table B.1). There was no effect of cover crop or date on stink bug abundance in field during either year, though boll injury has been suggested as more accurate indicator of stink bug pressure than total or mean abundance (Reay-Jones et al., 2010; Toews et al., 2010). A total of 2,639 bolls were assessed for stink bug feeding injury over 2 years. There was no effect of cover crop treatment on the proportion of cotton bolls injured in 2017 (GLMM: χ^2^_2, 31_ = 1.35; *p*=0.509). However, cover crop treatment significantly influenced the proportion of bolls injured in 2018 (GLMM: χ^2^_2, 43_ = 11.33; *p*=0.040; Fig. 3A). Overall stink bug pressure was much higher during the 2018 season than in 2017, and boll injury was significantly lower in rye plots compared to season conventionally managed no-cover plots (Fig. 3A).

**Fig. 3.**
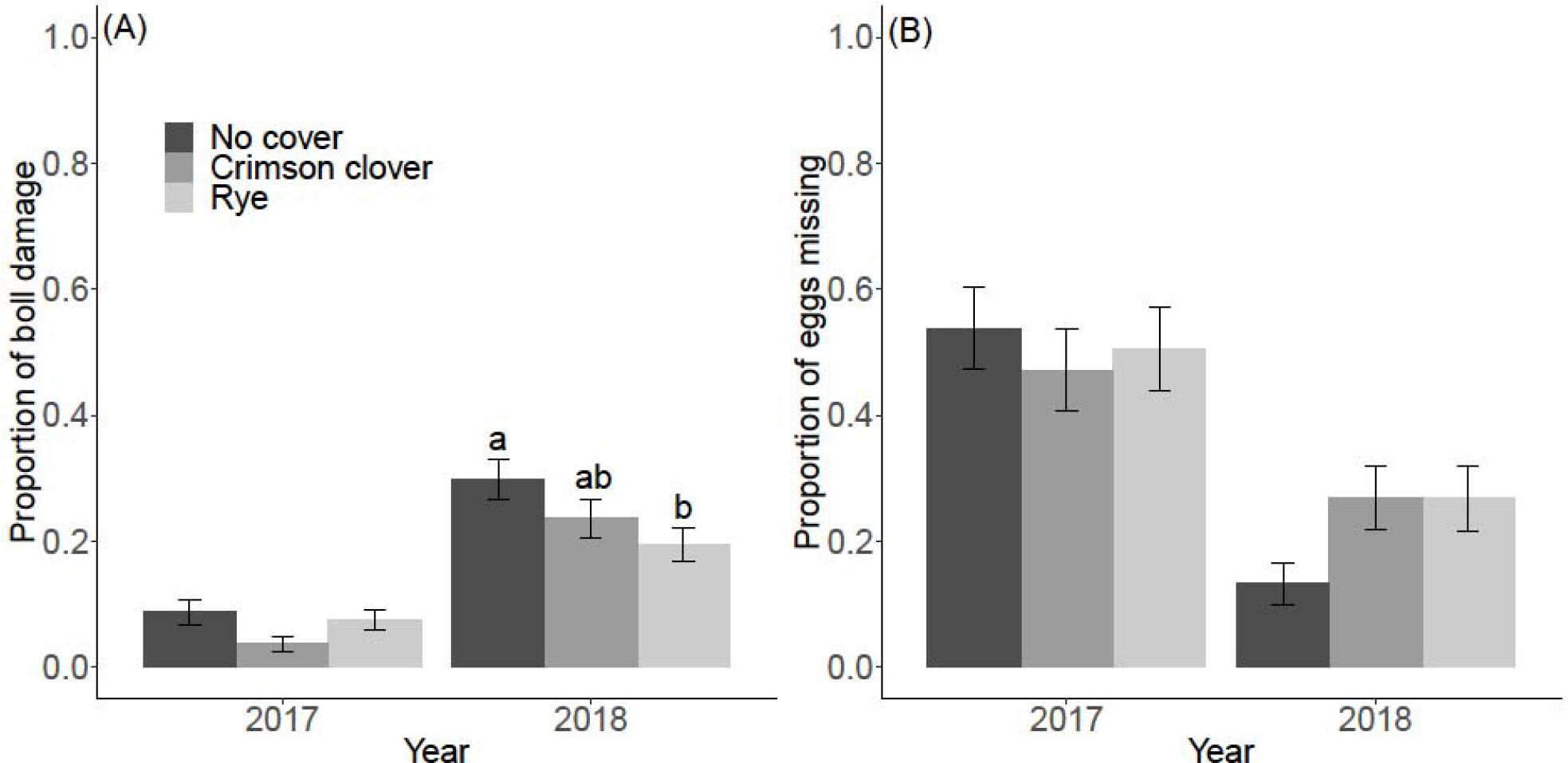
Boll injury from stink bug feeding (A) and *Nezara viridula* sentinel egg predation (B) shown for each cover crop treatment in 2017 and 2018. Y axes are proportion ranging between 0-1, with high proportion of bolls injured indicating higher pest pressure, and increased proportion of eggs missing indicating higher egg predation. Letters indicate significant differences among treatments in the same year (α<0.05).

### 3.2 Biological control services

Across both sample years, 336 *N. viridula* sentinel egg masses were placed in the field to estimate predation and parasitism rates. We found no effect of treatment on rates of egg predation in either 2017 (GLMM: χ^2^_2, 33_ = 1.35; *p*=0.51) or 2018 (GLMM: χ^2^_2, 45_ = 2.21; *p*=0.33). All emerging parasitoids were identified as *Trissolcus basalis*. Parasitism of southern green stink bug egg masses was very low for both sample years (0.05% in 2017 and 0.02% in 2018).

### 3.3 Cover crop biomass and cotton production

Cover crop biomass samples were used to evaluate the establishment and coverage of each cover crop treatment. Rye cover crops provided significantly higher initial biomass compared to crimson clover (Table 1). At the end of each growing season, yield and fiber quality were assessed to determine the effects of cover cropping treatments on cotton production. There was no effect of cover crop treatment on end of season cotton lint or seed yield alone, yet some aspects of cotton fiber quality were significantly influenced by cover crop treatment (Table 1; Supplementary Materials, Table C.2). Net return at both loan value and market value, including total cost of cover management, did not differ significantly among management treatments in 2017 (ANOVA_*loan*_: F_2,9_=0.93, *p*=0.430; ANOVA_*mkt*_: F_2,9_=0.29, *p*=0.753) or 2018 (ANOVA_*loan*_: F_2,9_=1.27, *p*=0.326; ANOVA_*mkt*_: F_2,9_=1.70, *p*=0.231; Fig. 4). Both loan value (ANOVA: F_1, 24_ = 54.80; p<0.0001) and market value (ANOVA: F_1,24_ = 8.62; p = 0.007) of cotton produced was greater in 2017 compared to 2018, independent of cover crop treatment (Fig. 4). For further details regarding cotton premiums and commodity value by treatment, see supplementary materials (Table C.3).

**Fig. 4.**
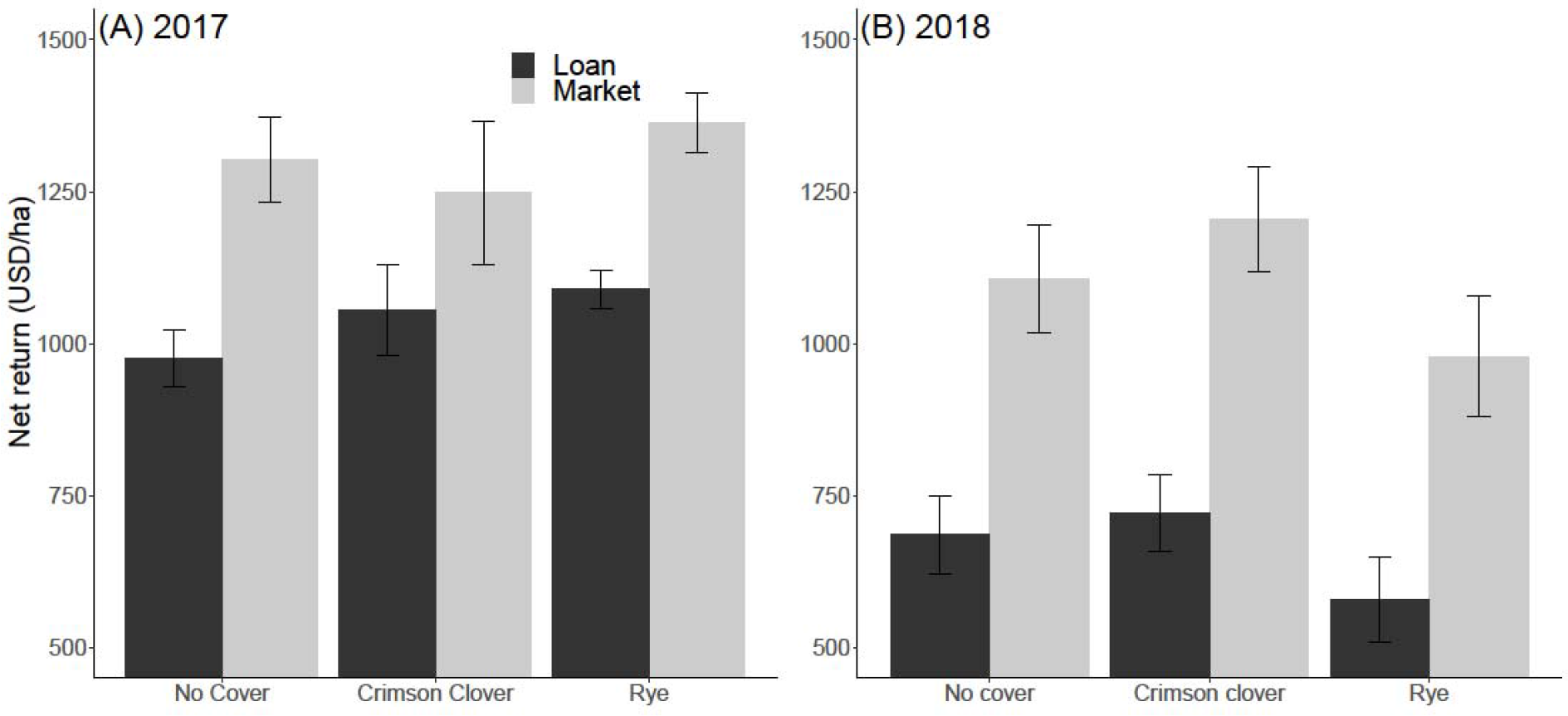
Mean net return on harvested cotton per hectare at both loan value and market value for each treatment in 2017 (A) and 2018 (B) with error bars showing SE of the mean. Net return (USD/ha) includes total production and management cost of cover crop establishment and termination, as well management costs associated with conventional cotton production with no cover crop.

## 4. Discussion

Winter cover crops provide multiple ecosystem services to agricultural systems (Hartwig & Ammon, 2002; Daryanto et al., 2018; Duzy & Kornecki, 2017; Adhikari et al., 2017). There is currently unharnessed potential of using cover crops to improve crop protection and reduce chemical inputs in annual production systems such as cotton. Our results supported aspects of our hypotheses by providing evidence of early season reduction in pest pressure, and correspondingly higher density and diversity of natural enemies. Importantly, our study demonstrates that integrating cover crops into the cotton production system results in similar costs of production and equivalent yields and quality of fiber produced.

Rye was more effective than either crimson clover or the no cover treatments at promoting higher natural enemy density and diversity. Differences in predator responses to cover crop treatments may be attributed to the type, and quality of resources provided by the cover crop, but also cover crop biomass. Quantity of cover crop residue influences weed suppression and other ecosystem services provided by cover crops (Toler et al., 2019; Finney et al., 2016; MacLaren et al., 2019). Thus, the significantly improved initial biomass of the rye cover crop (Table 1) may explain its increased effectiveness in promoting and harboring a high density and diversity of natural enemies. Increasing plant diversity is shown to increase the availability and diversity of herbivores (Siemann et al., 1998; Welti et al., 2017), which serve as prey when crop habitat is minimal in the early season (Moreira et al., 2016). The addition of habitat complexity to agricultural systems also correlates with prey consumption by generalist natural enemies through provisioning of alternative food sources (Staudacher et al., 2018). Our findings of increased natural enemy density in cover crops is consistent with previous literature focusing on predator response to habitat manipulations that improve the quality or quantity of habitat in an agroecosystem (Ribeiro & Gontijo, 2017; Depalo et al., 2017; Letourneau et al., 2011; Andow, 1991; Gurr et al., 2000; Holland et al., 2012). Therefore, we show that cover crops, especially rye, are an effective means of bolstering natural enemy communities ready to respond to pest immigration (i.e. early season thrips).

Cover crops provided notable suppression of early season thrips pests, with particularly low numbers of thrips nymphs found in rye in a majority of samples (Fig. 2B). Adult thrips are winged and mobile and their number and distribution in field indicate where thrips are present and active. Thrips nymphs are wingless and are unable to move between plots, therefore number of thrips nymphs is an effective estimate of thrips development and feeding pressure on seedling cotton. Our results suggest that the presence of a cover crop had a significant positive effect on early season thrips control compared to our conventional management treatment (Fig. 2), particularly the pressure and development of nymphal thrips on cotton. While we lack a direct test of the link between improved predator density and lower thrips counts, we did observe both higher numbers of predators and higher diversity, and lower numbers of thrips developing on cotton. Further research is needed to test proposed mechanisms and competing hypotheses for early season thrips response to cover crop use and habitat complexity (González-Chang et al., 2019; Brévault & Clouvel, 2019). Previous studies in cotton have found similarly reduced thrips pressure in cover cropped plots (Toews et al., 2010; Olson et al., 2006) suggesting that cover crops can provide a measure of thrips control without an in-furrow or foliar insecticide application.

Additionally, cover crops can provide relief from pest pressure into the late season, despite the similarities in the abundance and diversity of predators among treatments (Fig. 1). Our results indicate that when stink bug pressure is low, boll injury was similar across treatments. However, our results suggest that in years with high stink bug pressure, cover crops reduce levels of stink bug damage as much as 10% when compared to conventional management practices (Fig. 3A). Predation and parasitism of stink bug eggs has been shown provide a measure of control of pests in the field (Tillman, 2011), although we found no link between cover crop treatment and enhanced late season predator communities or the predation of *N. viridula* eggs (Figure 3B). Regardless of the exact mechanism, we show that cover crops can provide relief from high stink bug pressure, reducing the injury on developing bolls and potentially mitigating the impact on cotton production. Stink bug feeding and injury can have significant effects on cotton yield and quality, through direct damage of cotton seeds and lint resulting in abortion of young bolls, lint discoloration, and reduced lint production (Wene & Sheets, 1964; Barbour et al., 1988; Roach, 1988). Furthermore, feeding by *N. viridula* can result in the introduction of microorganisms resulting in boll rot (Willrich et al., 2004) or bacterial pathogens which affect boll development (Medrano et al., 2007), making evaluation of both end of season lint yield and quality important for determining the overall effect of cover crops and pest pressure on cotton production.

One of the primary concerns with the use of cover crops and strip tillage is the yield gap between conservation practices and conventionally managed cash crops (Ponisio et al., 2015; Reganold & Wachter, 2016; Tilman et al., 2011). Here we show production costs and net return when using cover crops are competitive with common conventional production methods. Production value calculations used in this study provide estimates for the value of cover crops on production cost and maintenance over two years, although several studies have highlighted additional long-term benefits of cover crop use (Reeves, 2018; Daryanto et al., 2018; Poeplau & Don, 2015). Some recent studies indicated that using cover crops may reduce inorganic fertilizer demand (Mahama et al., 2016; Plaza-Bonilla et al., 2017; Wilson et al., 2019), and in the current study we held all inputs constant across treatments. Further consideration for reduced pesticide applications for weeds and insect pests will enhance our estimates of overall costs.

In closing, our study indicates positive effects of cover crops on natural enemy communities, and possible applications of cover crops to improve biological control as part of integrated pest management programs. We highlight the seasonal aspect of cover cropping benefits on biodiversity and biological control, showing strong early season effects on predator communities and reductions of pest pressure into the late season. We provide evidence to support the use of cover crops to improve the sustainability of current agricultural systems by reducing the need for synthetic inputs while maintaining competitive levels of production and promoting local biodiversity. Although relationships between cover crops, density and diversity of predators, and pest pressure are evident, further research is needed in guiding pest management thresholds and inputs into the system in relation to cover cropping system used and quality of the cover crop. Lastly, research demonstrates that native predators are particularly important in cotton production (Naranjo 2018), and the effectiveness of predator diversity in providing biological control is closely linked to the composition and function of predators in the field (Heimpel & Mils, 2017); therefore, our future work will untangle how management strategies influence the composition and functional roles of native predators in cotton systems.

## Acknowledgements

Special thanks to Anthony Black for plot preparation and management. Additionally, the authors express thanks to Melissa Thompson, Zachary Wainwright, Abigail Borem, Emmalee Milner, Rachel Perez, Shereen Xavier, Lauren Perez, and Sarah Hobby for help with field collections and sample processing. This work was supported, in part, by the University of Georgia, USDA-NIFA Multistate Hatch Project GEO00884-S1073, Georgia Cotton Commission project 15-156GA, and Cotton Incorporated. Mention of trade names or commercial products in this publication is solely for the purpose of providing specific information and does not imply recommendation or endorsement by the University of Georgia.

