## Appendix A for "Cover crops improve early season natural enemy recruitment and pest management in cotton production"

Table A.1a) 2017 full plot management details including planting, irrigation, chemical pesticide and mineral fertilizer applications.

| Date | Management |
| --- | --- |
| 12/2/16 | Planted covers |
| 2/14/17 | 35units N/acre |
| 2/20/17 | Spread 1000lbs Lime |
| 3/29/17 | 1qt Glyphosate (Covers only) |
| 4/17/17 | Field Cultivated no-cover plots |
| 4/18/17 | 30-90-80 Truck Spread |
| 5/3/17 | Strip-tilled all plots |
| 5/3/17 | 1 qt Pendimethalin behind Striptill |
| 5/8/17 | Planted Cotton (PHY 490; 7.9 seeds/m) |
| 5/8/17 | Applied 10ozs Reflex, 10ozs Diuron, 1 qt Glyphosate |
| 5/8/17 | Irrigated .35 in |
| 5/11/17 | Irrigated .50 in |
| 5/16/17 | Irrigated .35 in |
| 5/18/17 | Irrigated .50 in |
| 5/30/17 | 29ozs Liberty |
| 6/17/17 | Irrigated .50 in |
| 6/26/17 | 0.1ozs Envoke, 1qt Glyphosate, 1qt/100 Surfactant |
| 6/28/17 | 70 units N sidedressed (all plots) |
| 6/29/17 | Irrigated .35 in |
| 7/11/17 | 1 qt MSMA, 1 qt Diuron, .10 oz Envoke Layby |
| 7/12/17 | Irrigated .75 in |
| 7/22/17 | Irrigated .75 in |
| 7/25/17 | 12ozs Mepiquat, .25 lb Boron |
| 7/27/17 | Irrigated .75 in |
| 7/28/17 | Irrigated .75 in |
| 8/1/17 | Irrigated .75 in |
| 8/6/17 | Irrigated .75 in |
| 8/15/17 | Irrigated .75 in |
| 8/20/17 | Irrigated .75 in |
| 8/24/17 | Irrigated .75 in |
| 9/1/17 | Irrigated .75 in |
| 9/4/17 | Irrigated .75 in |
| 10/3/17 | 4ozs Dropp, 42ozs Ethephon, 10ozs Def |
| 10/31/17 | Harvest |

Table A.1b) 2018 full plot management details including planting, irrigation, chemical pesticide and mineral fertilizer applications.

| Date | Management |
| --- | --- |
| 11/20/17 | Planted covers |
| 11/27/17 | 26 units N/acre |
| 11/27/17 | Irrigated 1.00 in |
| 1/22/18 | Spread 1000lbs Lime |
| 3/5/18 | 1st cut with disk (No-cover only) |
| 4/2/18 | 1qt Glyphosate (Covers only) |
| 4/3/18 | 2nd cut with disk (No-cover only) |
| 4/3/18 | 30-60-110 Truck Spread |
| 4/20/18 | Strip-tilled all plots |
| 4/28/18 | 1qt Grmoxone, 10ozs Reflex, 10ozs Diuron |
| 4/28/18 | Planted Cotton (PHY 440; 3.4 seed/ft) |
| 5/1/18 | Irrigated 0.35 in |
| 5/3/18 | Irrigated 0.50 in |
| 5/14/18 | 29 ozs Liberty, 2 pts Warrant |
| 6/9/18 | 1qt Glyphosate, 2ozs Staple |
| 6/9/18 | 65 units N side-dressed (all plots) |
| 6/9/18 | Irrigated .35 in |
| 6/11/18 | Irrigated .35 in |
| 6/23/18 | Irrigated .75 in |
| 6/27/18 | 1 qt MSMA, 1 Qt Diuron, 1 qt COC, .10 Oz Envoke |
| 7/9/18 | 12ozs Mepiquat, .25lbs Boron |
| 7/9/18 | Irrigated .75 in |
| 7/12/18 | Irrigated .75 in |
| 7/19/18 | Irrigated .75 in |
| 7/24/18 | Irrigated .75 in |
| 7/27/18 | Irrigated .75 in |
| 8/15/18 | Irrigated .75 in |
| 8/19/18 | Irrigated .75 in |
| 9/21/18 | 6ozs Folex, 4ozs Drop, 42ozs Ethephon, 1qt Glyphosate |
| 10/4/18 | Harvest |

Table A.2) Arthropod suction sample dates corresponding to each major cotton development stage.

| **Cotton stage** | **2017** | **2018** |
| --- | --- | --- |
| Seed | 5/16/17 | 5/8/18 |
| Seedling | 5/31/17 | 6/1/18 |
| Leafy growth | 6/23/17 | 6/19/18 |
| Squaring | 7/19/17 | 7/10/18 |
| Flowering | 8/2/17 | 7/26/18 |
| Boll Development | 8/14/17 | 9/5/18 |
