## Appendix B for "Cover crops improve early season natural enemy recruitment and pest management in cotton production"

Table B.1). Total number of stink bugs collected from sweep net samples in each treatment across all sample dates. Counts include both adults and identifiable nymphs.

|  | *2017* |  |  | *2018* |  |  |  |
| --- | --- | --- | --- | --- | --- | --- | --- |
| **Stink Bug Taxa** | **No cover** | **Crimson clover** | **Rye** | **No cover** | **Crimson clover** | **Rye** | **Total** |
| *Nezara viridula* | 13 | 4 | 4 | 11 | 0 | 12 | 44 |
| *Euschistus servus* | 1 | 0 | 0 | 6 | 5 | 5 | 17 |
| *Acrosternum hilare* | 0 | 0 | 0 | 1 | 1 | 1 | 3 |
| *Euschistus quadrator* | 0 | 0 | 0 | 1 | 0 | 0 | 1 |
| Other stink bugs | 0 | 2 | 1 | 7 | 6 | 3 | 19 |
| Total | 14 | 6 | 5 | 26 | 12 | 21 | 84 |

Table B.2) Total abundance of predators by taxa (family level or higher) in each treatment for 2017 and 2018. Listed in order by most abundant to least (combined 2017&2018).

|  | *2017* |  |  | *2018* |  |  |  |
| --- | --- | --- | --- | --- | --- | --- | --- |
| Predator Taxa | Con | CC | Rye | Con | CC | Rye | Total |
| Lycosidae | 20 | 20 | 74 | 30 | 71 | 180 | 395 |
| Anthocoridae | 126 | 86 | 68 | 32 | 19 | 27 | 358 |
| Coccinellidae | 33 | 38 | 24 | 53 | 55 | 53 | 256 |
| Linyphiidae | 7 | 4 | 26 | 10 | 40 | 137 | 224 |
| Theridiidae | 25 | 17 | 19 | 47 | 53 | 62 | 223 |
| Staphylinidae | 6 | 4 | 16 | 14 | 26 | 98 | 164 |
| Geocoridae | 9 | 33 | 53 | 12 | 15 | 19 | 141 |
| Carabidae | 19 | 37 | 25 | 5 | 18 | 16 | 120 |
| Salticidae | 10 | 13 | 9 | 24 | 34 | 18 | 108 |
| Neuroptera | 24 | 7 | 10 | 25 | 24 | 13 | 103 |
| Gnaphosidae | 2 | 7 | 24 | 1 | 6 | 42 | 82 |
| Araneidae | 17 | 9 | 19 | 10 | 7 | 15 | 77 |
| Nabidae | 19 | 8 | 23 | 5 | 7 | 11 | 73 |
| Elateridae | 10 | 15 | 6 | 12 | 13 | 12 | 68 |
| Thomisidae | 6 | 7 | 6 | 14 | 19 | 14 | 66 |
| Oxyopidae | 3 | 11 | 11 | 10 | 8 | 5 | 48 |
| Reduviidae | 5 | 9 | 13 | 6 | 5 | 2 | 40 |
| Chilopoda | NA | NA | NA | 2 | 12 | 21 | 35 |
| Tetragnathidae | 4 | 7 | 4 | 4 | 3 | 11 | 33 |
| Dolichopodidae | 1 | 1 | 1 | 5 | 7 | 9 | 24 |
| Dermaptera | 1 | 3 | 5 | 1 | 0 | 7 | 17 |
| Miturgidae | 0 | 0 | 0 | 0 | 1 | 4 | 5 |
| Pisauridae | 2 | 0 | 0 | 1 | 1 | 0 | 4 |
| Dictynidae | 0 | 0 | 1 | 0 | 0 | 2 | 3 |
| Podisus | 0 | 0 | 0 | 0 | 0 | 2 | 2 |
| Mimetidae | 0 | 0 | 0 | 1 | 0 | 1 | 2 |
| Anyphaenidae | 0 | 0 | 0 | 1 | 0 | 0 | 1 |
| Cubionidae | 1 | 0 | 0 | 0 | 0 | 0 | 1 |
| Hahniidae | 0 | 0 | 0 | 0 | 0 | 1 | 1 |
| Philodromidae | 0 | 0 | 0 | 0 | 1 | 0 | 1 |
| Total | 350 | 336 | 437 | 325 | 445 | 782 | 2675 |

Table B.2. Mean (±1SE) predator density (no./m^2^) (a) and diversity (H) (b) for each treatment during each major cotton development stage in 2017 and 2018. Results of linear contrasts on the interaction between date and cover crop treatment are indicated with letters where different letters indicate significant differences between treatments at a given cotton stage (α<0.05). Asterisks (*) on cotton stages indicate where predator abundance or diversity significantly differed among treatments.

| *a) Predator Density (no./m^2^)* | |  |  |
| --- | --- | --- | --- |
| Cotton Stage | No Cover | Crimson Clover | Rye |
| *2017* |  |  |  |
| Seed* | 0.81(0.41)^a^ | 3.5(0.80) ^ab^ | 7.0(0.89)^b^ |
| Seedling | 0.75(0.28) | 1.92(0.38) | 3.92(0.89) |
| Leafy Growth | 2.17(0.35) | 2.58(0.47) | 6.17(0.86) |
| Squaring | 5.0(0.74) | 4.33(0.95) | 4.0(0.94) |
| Flower | 8.42(1.05) | 5.17(0.83) | 7.08(0.91) |
| Boll Development | 11.75(0.85) | 9.58(1.69) | 7.0(0.98) |
| *2018* |  |  |  |
| Seed* | 0.67(0.36)^a^ | 3.33(1.42)^a^ | 9.67(1.93)^b^ |
| Seedling* | 1.92(0.47)^a^ | 3.92(0.82)^a^ | 21.75(2.55)^b^ |
| Leafy Growth | 2.42(0.61) | 2.25(0.60) | 5.58(1.10) |
| Squaring | 6.33(0.79) | 5.0(0.62) | 6.42(0.92) |
| Flower | 9.0(1.34) | 8.08(1.19) | 9.92(0.94) |
| Boll Development | 6.67(0.87) | 6.92(1.78) | 8.0(1.18) |

| *b) Predator Diversity (H)* | | | |  |
| --- | --- | --- | --- | --- |
| Cotton Stage | No Cover | Crimson Clover | | Rye |
| *2017* |  |  | |  |
| Seed* | 0.0(0.0)^a^ | 0.52(0.12) ^ab^ | | 1.15(0.17)^b^ |
| Seedling | 0.11(0.07) | 0.31(0.12) | | 0.65(0.16) |
| Leafy Growth* | 0.49(0.13)^a^ | 0.40(0.13)^a^ | | 1.25(0.08)^b^ |
| Squaring | 1.12(0.13) | 0.68(0.17) | | 0.85(0.15) |
| Flower | 1.42(0.11) | 1.15(0.19) | | 1.50(0.14) |
| Boll Development | 1.28(0.10) | 1.02(0.13) | | 1.25(0.12) |
| *2018* |  | |  |  |
| Seed* | 0.12(0.08)^a^ | | 0.59(0.16) ^ab^ | 0.92(0.14)^b^ |
| Seedling* | 0.61(0.19)^a^ | | 0.92(0.15)^ab^ | 1.67(0.09)^b^ |
| Leafy Growth | 0.65(0.19) | | 0.42(0.17) | 0.93(0.19) |
| Squaring | 1.45(0.16) | | 1.39(0.10) | 1.36(0.20) |
| Flower | 1.79(0.11) | | 1.56(0.18) | 1.75(0.06) |
| Boll Development | 1.50(0.16) | | 1.30(0.10) | 1.39(0.17) |
