## Appendix C for "Cover crops improve early season natural enemy recruitment and pest management in cotton production"

C.1). The cottonseed yield was estimated using a conversation ratio of 1.412 pounds of seed per pound of lint, which is obtained from the Upland Cotton Loan Calculator Program Decision Aid developed by Cotton Incorporated. Cottonseed prices in each year were obtained from U.S. Department of Agriculture National Agricultural Statistics Service. The prices for cotton lint include the cotton loan price and market price to compare the effect of prices on the profitability of different treatment. Cover crop seed costs come from the actual costs of the field experiment. Cover crops were planted using the costs for no-till drill (Lazarus 2017, 2018). The costs for two additional tandem disk with field cultivator were used for conventional tillage (Lazarus 2017, 2018). Cost for a strip-till rig for cover crop from UGA Cotton Budget (2017, 2018). The costs for row crop planter from Lazarus (2017, 2018) were used for planting cotton. Herbicide costs come from the actual costs of the field experiment and application cost for herbicide comes from Lazarus (2017, 2018), which is the total cost per acre of a self-propelled boom sprayer. Picking and moduling costs come from the Upland Cotton Loan Calculator Program Decision Aid developed by Cotton Incorporated. Ginning costs come from UGA Cotton Budget (2017, 2018). All costs include fuel, lubricants, repairs and maintenance, labor, electricity, depreciation (depreciation is both time-related and use related), and overhead costs (interest, insurance, and housing). Four trips of herbicide were used for conventional tillage, and five trips of herbicide application were used for the cover crop treatments.

Table C.2) Mean production values and standard errors (SE) for each cover crop treatment at loan value and market value for cotton crop in 2017 (a) and 2018 (b). Includes price premium based on fiber quality, price at loan and market value for cotton lint, and production cost used for net return ($/acre). The price premium for cotton fiber includes fiber grade and length, cotton micronaire (mic), uniformity, and extraneous matter premiums. Total cost includes the costs of planting cover crop and cotton, herbicide and application costs during the season, and costs associated with picking and moduling costs as well as ginning cost. All units are in USD per acre unless otherwise stated.

(a) 2017

| Cover | Value | Price premium (cents/lb) | Lint price (cents/lb) | Lint value | Seed value | Gross return | Total cost | Net return |
| --- | --- | --- | --- | --- | --- | --- | --- | --- |
| No cover | Loan | 3.81 (0.37) | 53.30(0.37) | 468.50(20.45) | 88.13(3.69) | 556.63(24.09) | 120.42(0) | 394.95(18.71) |
| Crimson clover | Loan | 4.56 (0.15) | 54.05(0.15) | 485.13(34.26) | 90.00(6.43) | 575.13(40.68) | 170.19(0) | 427.38(30.22) |
| Rye | Loan | 4.71 (0.25) | 54.20(0.25) | 499.63(14.04) | 92.38(2.47) | 592.00(16.49) | 144.39(0) | 441.13(12.75) |
| No cover | Market | 4.66(0.56) | 80.36(0.56) | 706.25(30.46) | 88.13(3.69) | 794.38(34.09) | 120.42(0) | 527.33(28.18) |
| Crimson clover | Market | 6.03(0.23) | 81.72(0.23) | 733.25(51.78) | 90.00(6.43) | 823.25(58.20) | 170.19(0) | 505.31(47.74) |
| Rye | Market | 6.11(0.34) | 81.81(0.34) | 754.25(21.10) | 92.38(2.47) | 846.63(23.55) | 144.39(0) | 551.36(19.80) |

(b) 2018

| Cover | Value | Price premium (cents/lb) | Lint price (cents/lb) | Lint value | Seed value | Gross return | Total cost | Net return |
| --- | --- | --- | --- | --- | --- | --- | --- | --- |
| No cover | Loan | 1.79(0.82) | 53.79(0.82) | 447.63(27.42) | 89.25(4.64) | 536.88(31.98) | 127.39(0) | 277.36(25.70) |
| Crimson clover | Loan | 3.41(0.58) | 55.41(0.58) | 510.88(26.92) | 99.00(5.15) | 609.88(31.97) | 171.96(0) | 291.79(25.28) |
| Rye | Loan | 3.82(0.29) | 55.82(0.29) | 423.00(31.04) | 81.25(5.96) | 504.25(37.00) | 149.16(0) | 234.47(28.17) |
| No cover | Market | 2.31(1.24) | 74.28(1.24) | 618.00(37.60) | 89.25(4.64) | 707.25(42.11) | 127.39(0) | 447.74(35.90) |
| Crimson clover | Market | 4.76(0.94) | 76.73(0.94) | 706.88(36.60) | 99.00(5.15) | 805.88(41.61) | 171.96(0) | 487.79(35.01) |
| Rye | Market | 5.16 (0.64) | 77.14(0.64) | 584.63(43.20) | 81.25(5.96) | 665.88(49.14) | 149.16(0) | 396.09(40.35) |

| **Fiber quality metric** | **Description** |
| --- | --- |
| Lint Yield | Weight of marketable cotton lint (after seed removal). Measured in both lbs/acre and kg/ha. |
| Color grade | gradations of grayness and yellowness in the cotton based on measurements of both Rd and +B |
| Staple | Length of cotton fibers in 32nds of an inch |
| Mic(micronaire) | Cotton’s fineness reported to the nearest tenth. Measures resistance to air flow per unit mass |
| Strength | The fiber strength measurement is made by clamping and breaking a bundle of fibers with a 1/8-inch spacing between the clamp jaws. Results are reported in terms of grams per tex to the nearest tenth. A tex unit is equal to the weight in grams of 1,000 meters of fiber. Therefore, the strength reported is the force in grams required to break a bundle of fibers one tex unit in size. |
| Rd | Gradation of grayness of cotton fiber measured as % reflectance. |
| +B | Yellowness of cotton fiber. Greater +B value indicated increasing yellowness. |
| HVI Length | Length of cotton fibers measured in inches. |
| Uniformity | Uniformity index is a measure of the degree of uniformity of the fibers in a sample to the nearest tenth. Displayed as a percentage with a higher percentage indicating higher uniformity cotton |

Table C.3. Descriptions of fiber quality metrics tested (USDA, 2004).
